# Tau Clearance Reverses Neuronal Dysfunction in Both Young and Aged *C. elegans*

**DOI:** 10.1101/2025.06.23.661052

**Authors:** T. Carroll, G.V.W. Johnson, K. Nehrke

## Abstract

Alzheimer’s Disease (AD) is an age-related dementia and presents a growing medical and economic burden as the average human lifespan continues to rise. AD is classically diagnosed via the accumulation and aggregation of two major proteins: amyloid-β and tau. To date, potential and FDA-approved therapies designed to clear these aggregates at best delay rather than prevent disease, indicating that the root cause of AD lay upstream of aggregate formation. Tau protein’s phosphorylation is critical for AD progression, and phosphorylation at Threonine 231 is thought to be an early disease-associated, “gatekeeper” event. Previously, we showed that genomic, single-copy insertion of phosphomimetic human tau (T231E) into *C. elegans* mechanosensory neurons induced age-dependent deficits in light-touch sensation. Herein, we have generated new *C. elegans* models which express pan-neuronal human tau to assess the idea of selective vulnerability and whether specific neuronal behaviors might be impacted preferentially. We also tested whether tau clearance via an Auxin Inducible Degron (AID) could reverse these deficits. Despite our hypothesis that prolonged stress in older animals would induce irreversible metabolic rewiring or maladaptation, tau depletion rescued known behavioral deficits at all ages tested, including in old worms which displayed the most overt phenotypes. Taken together, our data suggest that neuronal dysfunction induced by phosphorylated tau is reversible and provides reassurance that current early-phase therapeutic efforts aimed at reducing soluble tau levels in AD patients may prove effective.

## Background and Introduction

Alzheimer’s disease (AD) is classically and clinically defined by the accumulation of two insoluble protein aggregates: extracellular amyloid-β plaques and intraneuronal neurofibrillary tangles (NFTs) composed of aberrantly modified tau protein ^1^. Despite decades of intensive research, AD etiology is poorly understood, available therapies are scarce, and AD progression can be slowed but cannot be halted ^2^.

Elucidating the mechanisms that drive AD is thus paramount for developing effective treatments that alleviate AD burden. Many modern AD therapies aim to clear or degrade amyloid-β and tau, hoping to prevent their aggregation and thus halt disease progression. However, emerging evidence suggests that clearance-based strategies may not be as straightforward or effective as initially hoped. Aducanumab, an amyloid- β targeting immunotherapy, was the first FDA approved AD drug ^3^. However, applying the exclusion criteria associated with aducanumab significantly limits patient access despite its FDA approval, and even in eligible patients its effectiveness in slowing cognitive decline is still being researched. ^3^. Many other amyloid-β and tau targeting clearance strategies are currently in the pipeline, and while early results have claimed mixed success, none have proved to be the silver bullet the AD field had envisioned ^4, 5^. There could be many reasons that clearance strategies to date have proved ineffective, including a treatment window applied after overt neurodegeneration has already occurred, permanent cellular changes which aren’t reversed upon clearance, or the mis-targeting of epitopes which are downstream and simply correlative to the root cause of disease.

As AD progresses, tau protein undergoes many post-translational modifications (PTMs), and its pathologic contribution to AD may begin upstream of its oligomerization or aggregation. Tau accumulates PTMs as it transitions from soluble monomer to insoluble aggregate in AD, with phosphorylation being the earliest and most abundant modification ^6^. Early-disease phosphorylation events modulate tau conformations ^7, 8^, tau localization ^9, 10^ and tau oligomerization ^11^. Phosphorylated threonine 231 (pT231) is one of the earliest tau PTMs to enrich in AD ^6, 12^, heralds subsequent NFT formation where it enriches ^13^, and is a promising biomarker for AD diagnosis well before aggregate formation ^14^. Furthermore, studies in post-mortem human tissue suggest that tau PTMs accumulate in a stereotypical pattern as sporadic AD progresses – highlighting the potential for pT231 as an early marker for disease inception ^6^. In line with this, our previous work in *C. elegans* demonstrated that single-copy expression of T231E drives functional deficits in neurons in the absence of apparent aggregates and suppresses stress-induced mitophagy, which is not observed in strains expressing wild-type human tau^15^.

Herein, we report generating a novel transgenic *C. elegans* which pan-neuronally expresses the pathologic human tau variant (T231E), use it to explore whether T231E tau preferentially impacts certain behaviors, and address whether clearance can rescue observed phenotypes. By using Mos-mediated Single Copy Insertion to generate integrated, single-copy transgenes, we have focused on expressing tau at physiologically relevant levels and attempted to avoid confounds related to overexpression of an aggregation prone protein. Moreover, tau expression in our strains is driven by the Ultra Pan-Neuronal (UPN) promoter, a synthetic promoter designed to express equally in all neurons ^16^, which facilitates defining selective vulnerability to T231E tau among the 302 well characterized neurons of the adult hermaphrodite nervous system. Surprisingly, while our results show that pan-neuronal expression of an early-disease, pathologic tau variant leads to deficits within the touch neurons of *C. elegans*, as previously reported, there are relatively few other behaviors that are significantly impacted, and those only subtly. Even more surprising was the finding that depleting the toxic T231E tau can completely restore the touch-deficit phenotype at any age tested, including in older worms. Although further study will be necessary to determine the molecular mechanisms behind this transient disruption of mechanosensory neuron function, these initial results provide reassurance that emerging therapeutic strategies designed to eliminate soluble tau may prove to be effective at treating AD.

## Materials and Methods

### C. elegans Growth and Maintenance

Nematodes were maintained at 20°C on Nematode Growth Media (NGM) plates and seeded with live *E. coli* strain OP50, cultured overnight at 37°C at 220 rpm, as previously described ^17, 18^. For experimental assays, after synchronization by standard procedure with sodium hypochlorite, 4th larval stage (L4) hermaphrodites were selected and moved to test plates. The day after moving was considered adult Day 1. Animals were transferred daily to avoid populations of mixed age.

### Plasmid Construction

Briefly, individual elements and coding sequences for each sub-component of pUPN::AID::eGFP::tbb-2 3’UTR and pUPN::AID::eGFP::TauT4::tbb-2 3’UTR constructs were amplified using primers with overlapping homology and then stitched together using overlap extension PCR cloning (Table 1) ^19^. pTC4 codes for human tau tagged with a minimal degron sequence in a pFH6.II *C. elegans* expression vector that includes GFP, synthetic introns, and a 3’ untranslated region. PCR sewing products were inserted into pCFJ151 (Addgene, Watertown, MA) using traditional restriction-site cloning to generate MosSCI vectors for insertion at the ttTi5605 Ch. II site ^20^. Final plasmids, pTC10 (AID::eGFP) and pTC11 (AID::eGFP::TauT4), were sequenced completely (Plasmidsaurus, San Francisco, CA, USA).

**Table 1.**
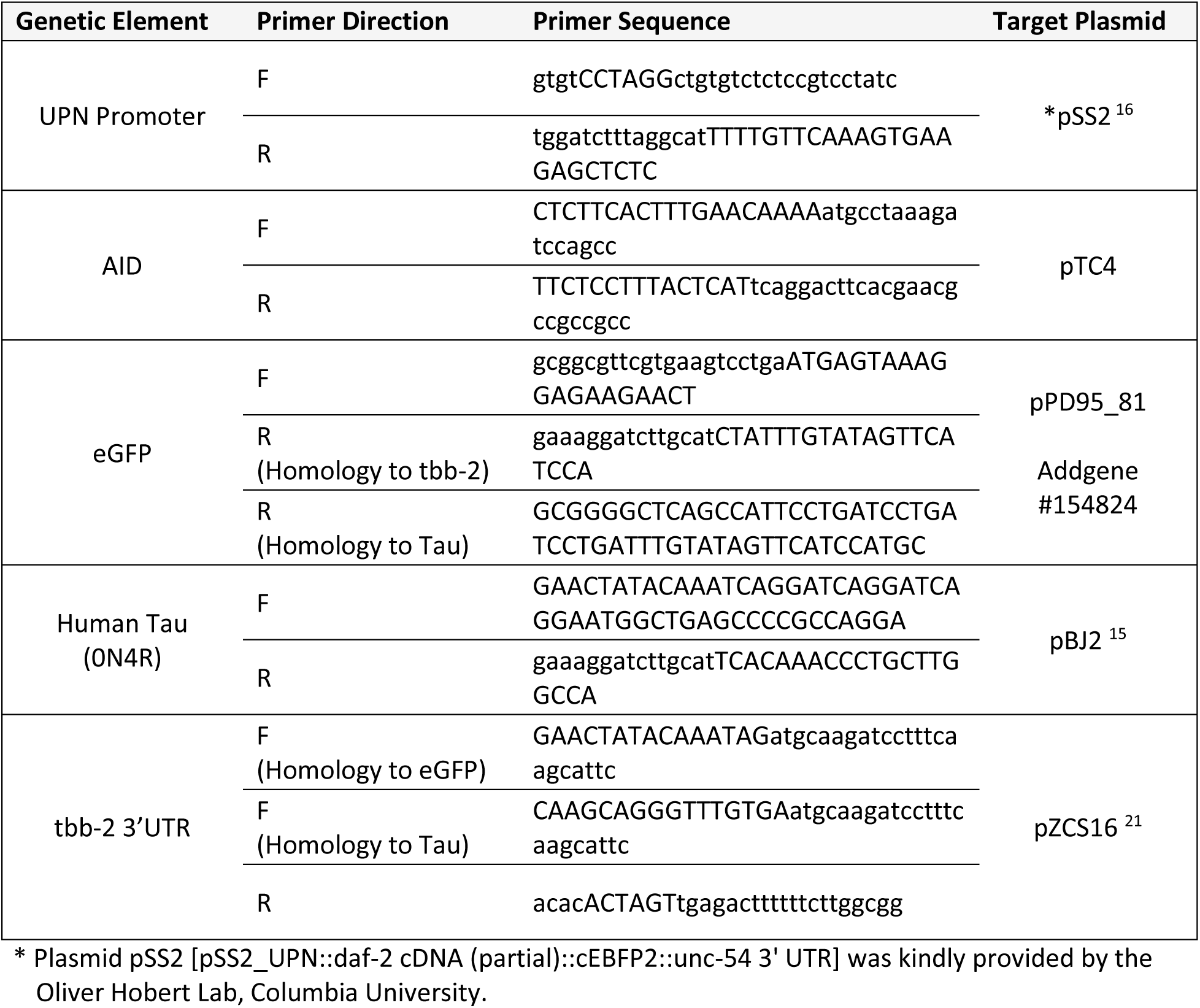
- Primers Used to Amplify Genetic Sub-elements for Overlap Extension PCR

### C. elegans Strain Generation

For all strains herein (see Table 2), the wild-type background strain is Bristol-N2 and all lab-generated and externally acquired strains were outcrossed to Bristol-N2 at least four times. Single-copy, pan-neuronal transgenic strains were created by integrating pTC10 and pTC11 into Ch. II site ttTi5605 using Mos-mediated Single-Copy Insertion ^20^. Both strains were sequenced completely through the insertion site. CRISPR-Cas9 gene editing was used to introduce site-specific mutations into the rnySi58 tau coding region via a *dpy-10* co-CRISPR strategy and oligonucleotide-mediated HDR using purified Cas9 RNP injection ^22,23^. Targeting crRNAs were from Dharmacon (GE Healthcare Dharmacon, Lafayette, CO) and were complexed to scaffolding RNAs for Cas9, with genomic recognition sites as follows: Tau T231 targeting RNA, 5’ACGGCGACTTGGGTGGAGTA’3 Tau T231E ssODN, 5’GTCCCTTCCAACCCCACCCACCCGGGAGCCCAAGAAGGTGGCCGTGGTCAGAGAGCCACCCAAGTCGCCGTCTTCCGCCAAGAGCCGCCTGCAGA’3 Strains were selected via pan-neuronal eGFP fluorescence, homozygosed, confirmed for successful gene insertion via genomic PCR with the following primers, and then outcrossed 4x to N2 to remove the co- CRISPR *dpy-10* mutant marker.

**Table 2.**
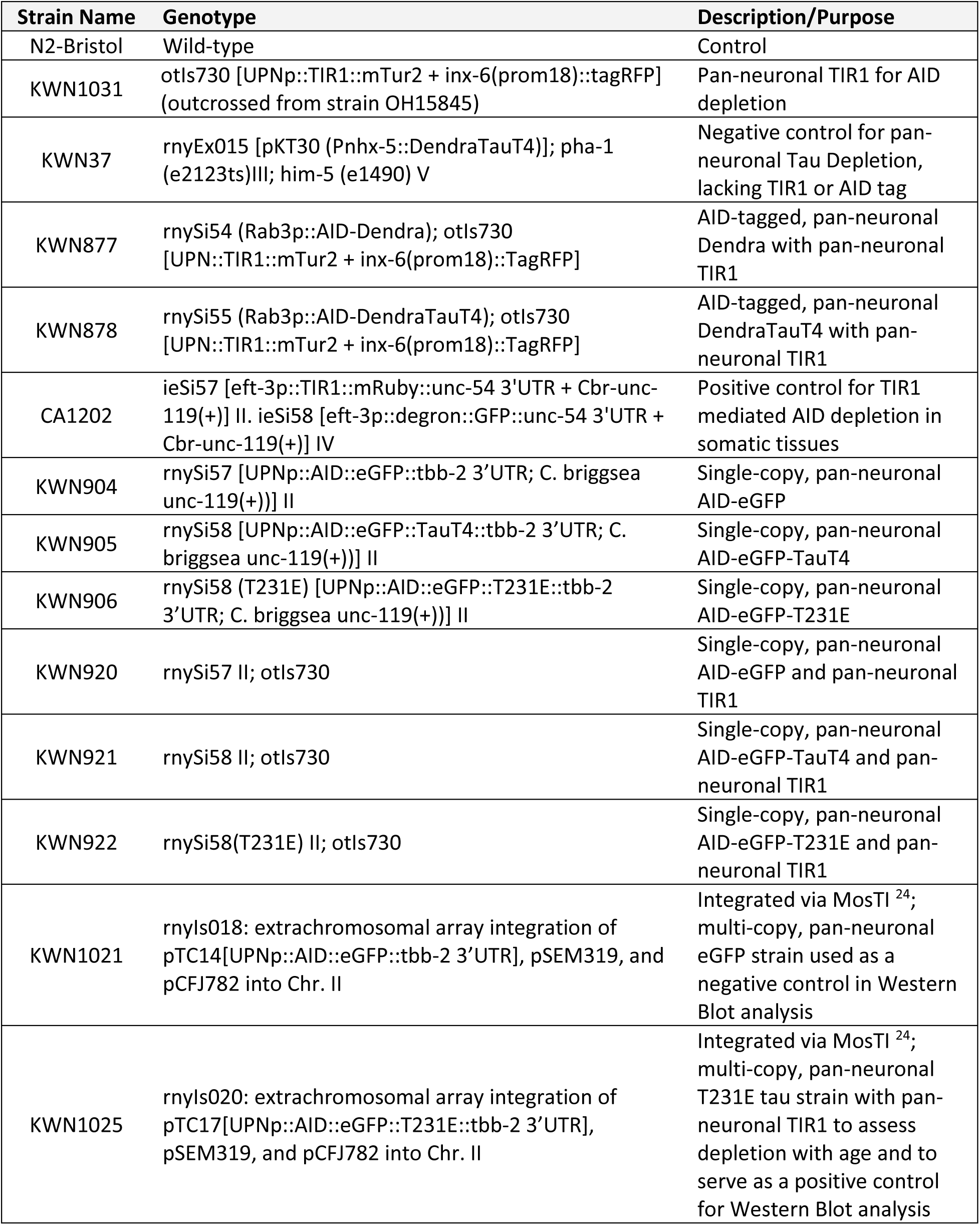
- *C. elegans* Strains Used in This Work

MosSCI ttTi5605-F, 5’GTTTTTGATTGCGTGCGTTA3’ MosSCI ttTi5605-R, 5’ACATGCTTCGTGCAAAACAG3’ MosSCI ttTi5605 insert-F, 5’CATCCCGGTTTCTGTCAAAT3’ Tau mutations were confirmed using the following genotyping primers: Tau geno-F2 5’ATCAGGGGATCGCAGCGG3’ Tau geno-R2 5’TGGTTTATGATGGATGTTGCCT3’

### Confocal Imaging

For pan-neuronal imaging, Day 1 animals were mounted on 2% agarose pads on glass slides and immobilized with 1 mM tetramisole hydrochloride (#L-9756; Sigma-Aldrich, St. Louis, MO). Imaging was performed using a Laser Scanning Confocal microscope (Inverted Leica DMi8 and LAX software). All images were acquired under the same exposure conditions with a 20x/0.75 objective (Air – HC PL Apochromat C52), and all genotypes were represented and imaged the same day. For clarity, autofluorescence was manually identified and pseudo-colored red in each image.

### AID Depletion and Validation

Age-synchronized animals were transferred to either regular NGM plates or NGM plates containing 4mM indole-3-acetic acid (IAA, A10556; Thermo Scientific Chemicals, Haverhill, MA) and left to deplete for 24 hours before assessing their protein content via fluorescent intensity. Imaging was performed using a Nikon Eclipse inverted microscope coupled to a six channel LED light source (Intelligent Imaging Innovation, Denver, CO), an ORCA-Flash4.0 V2 Digital CMOS camera (Hamamatsu Photonics, Bridgewater Township, NJ) and Slidebook6 software (Intelligent Imaging Innovation, Denver, CO). All images were acquired under the same exposure conditions and each experiment was imaged in one session. A depletion time course was established by taking Day 1 animals of a highly expressing, integrated tau strain containing pan-neuronal TIR1 (KWN1025) and imaging 1, 2, 4, and 24 hours after being transferred to 4mM IAA plates.

### Western Blot

Animals were maintained on 60mm NGM plates until nearly starved and collected using ice-cold M9 and washed 3x. The supernatant was discarded, and worm pellets were resuspended in 1x radioimmunoprecipitation assay (RIPA) buffer with 0.1% sodium dodecyl sulfate (SDS) and supplemented with protease and phosphatase inhibitors: 10 µM Bestatin (#200484; Calbiochem, San Diego, CA), 15 µM Pepstatin (#AG-CP3-7001; AdipoGen Life Sciences, San Diego, CA), 10µg/mL Leupeptin (#AG-CP3-7000M025; AdipoGen), 2µg/mL Aprotinin (#9087-70-1; MilliporeSigma, Burlington, MA), 1 mM PMSF (#837091, Roche, Basel, Switzerland), and 1x Halt^TM^ phosphatase inhibitor cocktail (#XB340823; Thermo-Fisher Scientific). Samples were freeze-cracked using liquid nitrogen three times, sonicated using a micro-tip at max amplitude for 20s two times (#S-3000; MISONIX Inc, Farmingdale, NY ), and then centrifuged at 4°C for 10min at 16,000g. Protein content was quantified using a bicinchoninic acid (BCA) assay, and 10µg protein was loaded into 4-15% gradient polyacrylamide gels (#4561083; Bio-Rad Laboratories, Hercules, CA) for separation then transferred to a nitrocellulose membrane. The membrane was blocked using 5% non-fat milk in Tris-buffered saline containing 0.05% Tween 20 (TBS-T). The membrane was probed using a 1:150,000 dilution of Dako rabbit, anti-tau antibody (1° - #A0024; Agilent, Santa Clara, CA)) overnight, washed three times using TBS-T, and then probed using a 1:5000 dilution of horse radish peroxidase (HRP) conjugated anti-rabbit antibody (2° - #R1006; Kindle Biosciences, Greenwich, CT) for 1 hour. Bands were developed using chemiluminescence (211595, Millipore, Immobilon Crescendo) and imaged using a KwikQuant Imager (Kindle Biosciences).

### Touch Sensitivity Assay

The behavioral response to being touched by an eyelash was adapted from an assay previously described ^25^. The animals were touched anteriorly specifically behind the terminal bulb of the pharynx with the eyelash, 10 times per animal, with a 10s gap between each touch. Typically, if the animal demonstrates an omega turn or if it reversed its direction after an anterior touch, the animal was scored as giving a positive response. Touch response percentage was generated by the number of times an animal responded to the touch stimulus over the total number of times they were touched. All touch response assays reported here were blinded by an independent researcher before experimentation.

#### Thrashing Assays

A drop of 2% agarose (#BP164-500; Thermo Fisher Scientific) was poured over a glass slide and allowed to dry and then 20 μl of M9 was poured on it. Age-synchronized animals were picked to that drop of M9 buffer. After 2 min in M9, thrashing rates were counted and recorded manually. A single thrash was defined as a complete change in the direction of the body down the midline. Animals that were motionless for 10 s were discarded from the analysis.

### Life-Span Assays

Embryos were obtained through conventional alkaline-hypochlorite treatment and placed on freshly grown OP50 seeded NGM plates. Fifteen animals from the 4th larval stage (L4) were transferred to individual 35 mm seeded NGM plates with a total 3 plates for each genotype. Each day they were transferred to new plates to avoid mixing of populations until they stopped producing offspring.

Simultaneously the worms were counted alive visually or with gentle prodding on the head. Animals were censored in the event of internal hatching of larva, body rupture, or crawling off the plate. The experiment was conducted at 20°C and scored until all the worms died ^26^.

### Locomotion Assays

Approximately 10 animals were transferred to NGM-agar plates and allowed to acclimate for 30 minutes before recording their movement for 10 minutes at 15 FP using a Nikon SMZ800 dissecting microscope equipped with Diagnostic Instruments Camera HRP042-CMT and XIMEA CamTool v4.28 capture software. Experiments examining locomotory speed (BLPS) were analyzed using the ImageJ plugin WrmTrck ^27^. Holistic locomotory experiments were conducted using Tierpsy Tracker to skeletonize and track more than 4,000 different aspects of worm locomotion, as described ^28^.

### Heatshock Assays

Twenty Day 1 animals were placed on NGM-agar plates, which were then covered with parafilm and placed in a 37°C incubator for 1 hour and 15 minutes ^29^. The animals were then visually scored as Normal, Paralyzed, or Dead after prodding with a platinum wire.

### PQT Stress Assays

Lifespan of animals experiencing chronic ROS stress induced by paraquat (PQT, #227320010, Thermo Scientific) exposure was assessed by moving Day 1 animals to plates containing 8mM PQT, adapted from^30^. Animals were transferred daily to avoid mixed populations, and they were scored as alive or dead via visual examination and gentle prodding on the head with a platinum wire.

### Associative Memory

Briefly, for each experimental strain, Day 1 animals were washed in M9 and transferred to conditioning NGM-agar plates (without food) in the presence or absence of isoamyl alcohol (IA, #AX1440-1, EM Science, Gibbstown, NJ) and left to starve for 90 minutes. Conditioned animals were then washed with M9 and transferred to assay plates to assess their chemotactic attraction or aversion to isoamyl alcohol, as previously described ^31^. The chemotactic index is calculated as (IA - T)/ (IA + T + S), where IA equals the number of worms residing in the quadrant containing IA, T equals the number of worms residing in the quadrant opposite the IA quadrant, and S equals the number of worms within the starting quadrant, at the end of the assay period.

### Statistical Analysis

All statistical analyses were conducted using Prism 8.0 (GraphPad Software Inc, Boston, MA), with alpha-error level of p < 0.05 considered to be significant. Data were averaged and represented as mean ± standard error (mean ± SEM) or as mean ± standard deviation (mean ± SD), depending upon the number of experimental replicates. In general, differences between matched pairs of samples were analyzed using student t-tests, and group differences were analyzed with either one-way or two-way ANOVA, depending upon the variables. For all tests herein, *P < 0.05, **P < 0.01, ***P < 0.001, ****P < 0.0001.

## Results

### Pan-Neuronal, Single-copy Human Tau Phosphomimetic Strains Possess Selective Deficits

Transgenic strains expressing Tau have historically utilized extrachromosomal arrays which can be stabilized by integration into the genome. While these strains have proven useful models ^32^, they do come with caveats regarding tau expression level, uniformity among tissues, and potential synthetic phenotypes due to disruption of endogenous genes during integration ^33, 34^. Previously, we showed that human tau expressed specifically in the mechanosensory neurons of *C. elegans* from single-copy gene insertions into a genomic safe-harbor loci could produce subtle, yet consistent, neuronal dysfunction that was selective to a phosphomimetic T231E AD associated mutation ^15^. Here, we expanded on this previous approach by generating a pan-neuronal equivalent to assess whether dysfunction occurs in other neuronal subtypes and if so, which subtypes are selectively vulnerable. In brief, the Ultra Pan-Neuronal (UPN) Promoter, an artificial, chimeric promoter comprised of regulatory sub-elements from *unc-11*, *rgef-1*, *ehs-1*, and *ric-19* ^16^, was used to drive equal levels of tau expression in all 302 neurons of the adult hermaphrodite. The primary expression construct consisted of 0N4R human tau ^35^, translationally fused to an eGFP fluorescent reporter to facilitate visualization, and an Auxin Inducible Degron (AID) tag for inducible degradation ^36^, inserted into a genomic safe-harbor loci using MosSCI (**Fig 1**) ^20^ . Like our previous model, T231E tau was generated via CRISPR-Cas9 gene editing of the transgenic cassette. An additional control strain was also generated, adopting an identical transgenic/genomic architecture to express AID::eGFP (**Fig 1**)

**Figure 1.**
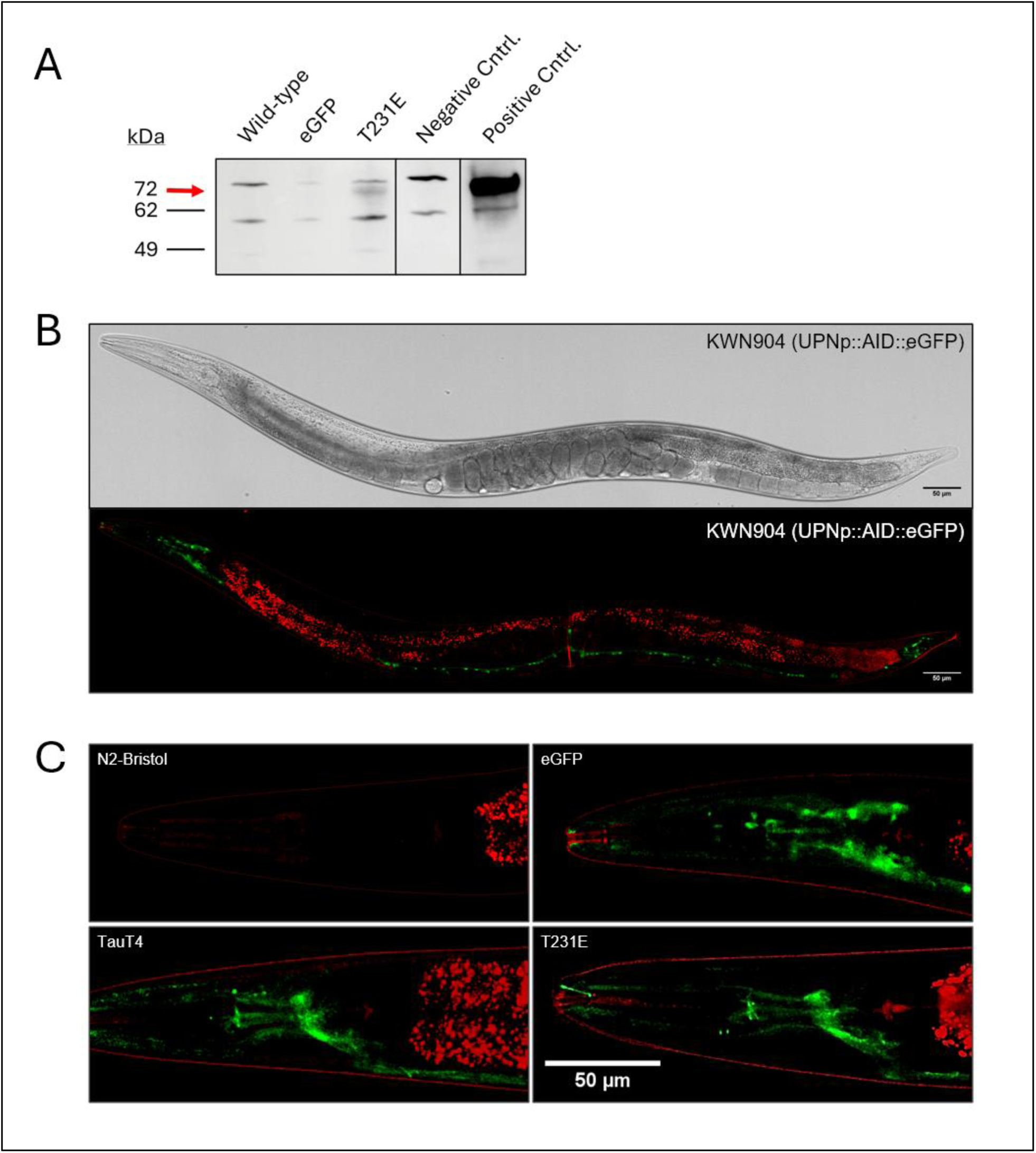
– Expression Pattern Characterization for Novel, Pan-Neuronal Tau Models (A) Western blot demonstrating measurable tau expression in the single-copy AID::eGFP::T231E animals. 10µg of protein lysate from wildtype (N2-Bristol), eGFP (KWN904, a single copy strain), and T231E (KWN906, a single copy strain) homogenates were separated on a 4-15% gradient polyacrylamide gel and transferred to a nitrocellulose membrane for western analysis. In addition, conventional transgenic strains were run as additional negative (KWN1021 for eGFP) and positive (KWN1025 for T231E Tau) controls. The red arrow indicates the specific band at the expected size of 72kDa. All other bands are non-specific. All samples were run on the same gel, transferred to the same membrane, and probed using a primary antibody for total tau. All lanes were exposed for the same amount of time, and the vertical lines denote where intervening lanes were removed for clarity. Extended blots can be found in Supplemental Figure 1. (B) Confocal images demonstrating the pan-neuronal expression of eGFP in strain KWN904 under the UPN promoter, a chimeric promoter specifically designed to drive equal levels of expression in all neurons ^16^. Autofluorescence has been manually pseudo-colored red for clarity. Scale bars = 50 µM. (C) Confocal images for N2 and pan-neuronal AID::eGFP (KWN904), AID::eGFP::TauT4 (KWN905), and AID::eGFP::T231E (KWN906) containing strains highlighting the expression pattern within *C. elegans* head neurons. In accordance with tau’s affinity for microtubules, the fluorescence for both AID::eGFP::TauT4 and AID::eGFP::T231E is predominantly localized to the nerve ring – a bundle of processes derived from the nearby head ganglia that wraps around the pharynx of the animal.

The single-copy pan-neuronal strains were imaged via confocal microscopy to confirm ubiquitous neuronal expression (**Fig 1B-C**). Interestingly, while eGFP was localized indiscriminately to the processes, soma, and even the nucleus, as has been observed previously in *C. elegans* ^37^, tau localization was more prominent in the processes, with the most visible fluorescent feature in both the TauT4 and T231E tau strains being the nerve ring, where many neuronal processes converge (**Fig 1C**). In addition, we confirmed that the expression level of the transgenes was approximately equivalent between cells, as expected from use of the UPN promoter (**Fig 1**). Finally, western blot analysis confirmed the expression of a polypeptide unique to the transgenic strains of the expected size of 72kDa for the fusion proteins (**Fig 1A**).

Behaviorally, pan-neuronal expression of T231E tau elicited a touch deficit (**Fig 2A**), like that reported previously in strains where tau expression was limited to mechanosensory neurons ^15, 38^. In contrast to previous studies, we also observed a small but significant deficit in strains expressing wildtype TauT4 when compared to non-transgenic wild type animals (N2-Bristol) (**Fig 2A**). This suggests that a palpable deficit requires its expression in non-mechanosensory neurons within the circuit, though whether these are independent or additive to its expression in mechanosensory neurons is unclear.

**Figure 2.**
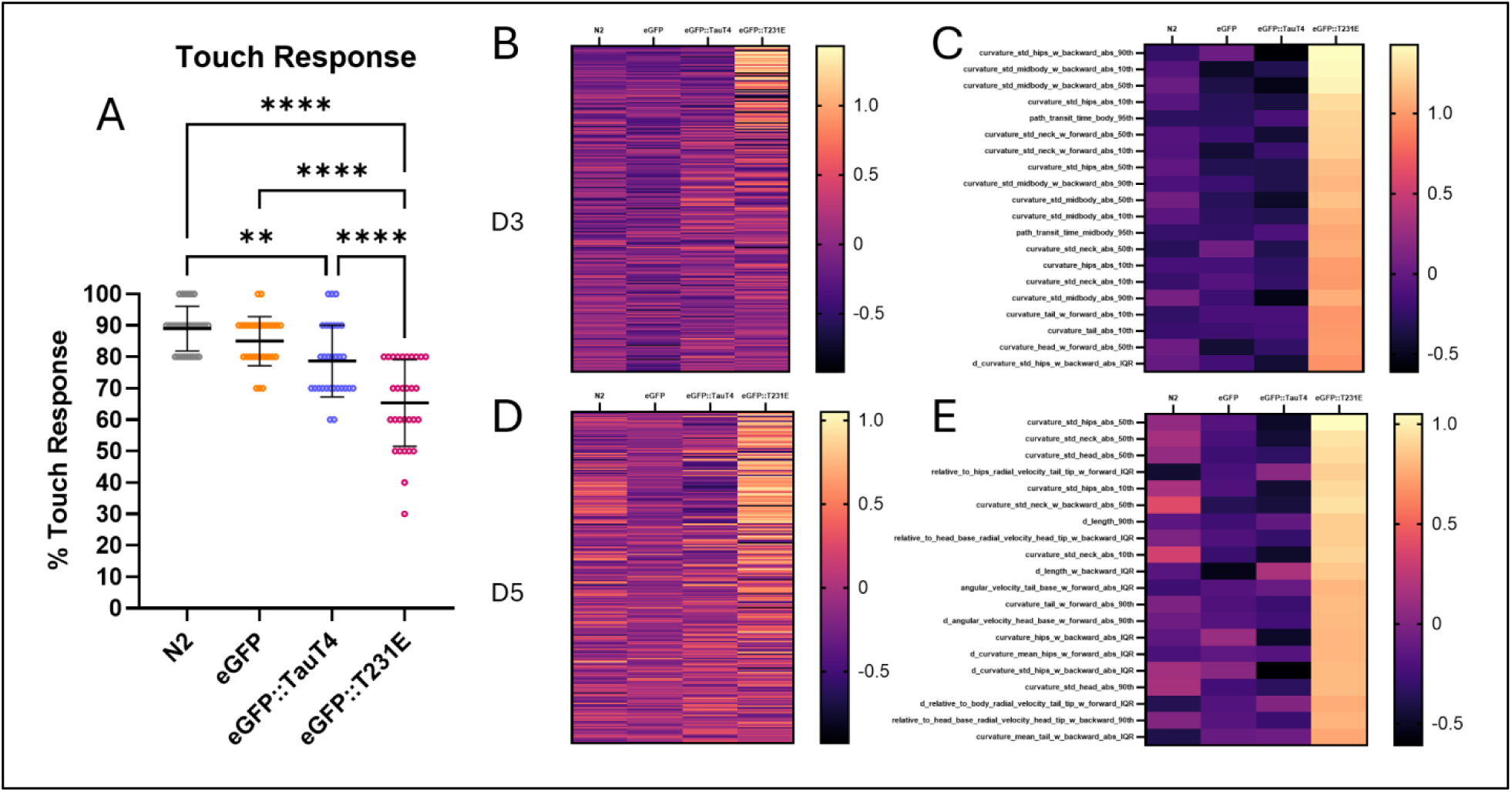
Touch-Sensation and Body Curvature are Impacted in T231E Tau Animals (A) Quantification of touch responsiveness in Day 5 adult N2 controls and animals expressing pan- neuronal eGFP, eGFP::TauT4, or eGFP::T231E. Data were calculated as percent responsiveness following ten repetitive light touches to the anterior body, and are plotted with the mean ± SD. Each point represents an individual animal, N = 30 animals from three biological replicates. Statistical analysis was performed using a one-way ANOVA followed by Tukey-corrected multiple comparisons. (B and D) An unbiased, clustering heat-map comparing locomotory features within the Tierpsy_256 library at day 3 (B) and day 5 (D). The Tierpsy_256 library is a hand curated list of the features generated through Tierpsy analysis that reduces the number of redundant comparisons between samples and is tailored to wholesale exploration of phenotypic differences in locomotion ^39^. Reported values are z-scores calculated using the grand mean and standard deviation of all samples, including controls. Initial analysis indicated that approximately a quarter of Tierpsy_256 features are different within the eGFP::T231E samples. (C and E) A heatmap focusing on the 20 features from (B) and (D), respectively, where the absolute eGFP::T231E z-score was highest, thereby highlighting the features where mimetic animals differ most. Nearly all these features relate to animal curvature on both days.

Although the transgenic strains lacked overt physiologic or behavioral abnormalities, T231E tau worms were still identifiable through blinded visual assessment by an experienced *C. elegans* experimentalist (data not shown), indicating that their locomotion was abnormal, albeit subtly and in a way that was not readily apparent. Hence, to obtain a robust and holistic picture of animal locomotion, we used Tierpsy Tracker, a complex worm tracking algorithm that measures and extracts 4,083 different features from videos of *C. elegans* foraging on solid media ^28, 39^. These features are categorized into ones that measure the morphology, path attributes, posture and curvature, as well as speed and velocity of the whole animal and individual body parts. Since Tierpsy’s inception, feature libraries have been established to highlight which measurements are most useful for determining phenotypic differences between animals. One such library is referred to as Tierpsy_256 and contains the top 256 most informative features for comparing worm locomotion ^28^. Tierpsy_256 analysis of our pan-neuronal T231E tau animals demonstrated that approximately a quarter of the features in the Tierpsy_256 library were strikingly different compared to controls at adult Days 3 and 5 (**Fig 2B and 2D**). By focusing the heatmap on the top 20 terms, it is apparent that nearly all of these relate to the posture, or curvature of the animal (**Fig 2C and 2E**). For example, the standard deviation of the curvature was different among all body parts for the T231E tau animals, indicating that the amplitude of their curvature was more erratic than that of the controls. Interestingly, this phenotype was limited to the T231E tau worms and was not apparent in the wildtype TauT4 expressing strain.

Next, we assessed other outputs, expecting to find additional, albeit perhaps subtle, impacts of T231E tau being expressed in every neuron. To our surprise, we were unable to identify such a phenotype. A selection of negative results is presented in **Supplemental Figure S1** including: lifespan (**Supplemental Fig S1A**), thrashing rates in liquid and basal locomotor speed on solid media, both of which are commonly reported phenotypes in pan-neuronal *C. elegans* tau models ^32^ (**Supplemental Figure S1D and S1E**), chemotaxis learning, which is a form of associative memory ^40, 41^ (**Supplemental Fig S1D**), and stress tolerance, including to acute heat and chronic exposure to paraquat (PQT), a redox cycler that generates ROS (**Supplemental Figure S1B and S1C**) ^30^. While these negative results do not preclude a role for tau in mediating other behaviors, the apparent selectivity may suggest that less nuanced effects are limited to mechanosensory touch neurons, which have remarkably long processes, as explored below in the Discussion.

### AID-tagged Tau Depletion Rescues Touch Deficits at All Ages

To determine whether our phosphomimetic tau creates irreversible dysfunction with age, we next asked whether T231E depletion could suppress our most robust phenotype: the touch deficit. Briefly, the Auxin Inducible Degron (AID) system requires three major components to mediate depletion of a select protein: 1) a protein labeled with an AID tag which binds auxin, a naturally occurring plant hormone, 2) a TIR1 protein complex which also binds auxin and poly-ubiquitinates the AID tagged protein and targets it for degradation by the proteasome, and 3) exogenous auxin supplementation to bring the AID tag and TIR1 complexes together ^36^ (**Fig 3A**). In our case, the transgenic proteins were labeled with AID, TIR1 was expressed from a pan-neuronal integrated transgene (*otIs730*) ^42^, and NGM plates were supplemented with a natural auxin (IAA).

**Figure 3.**
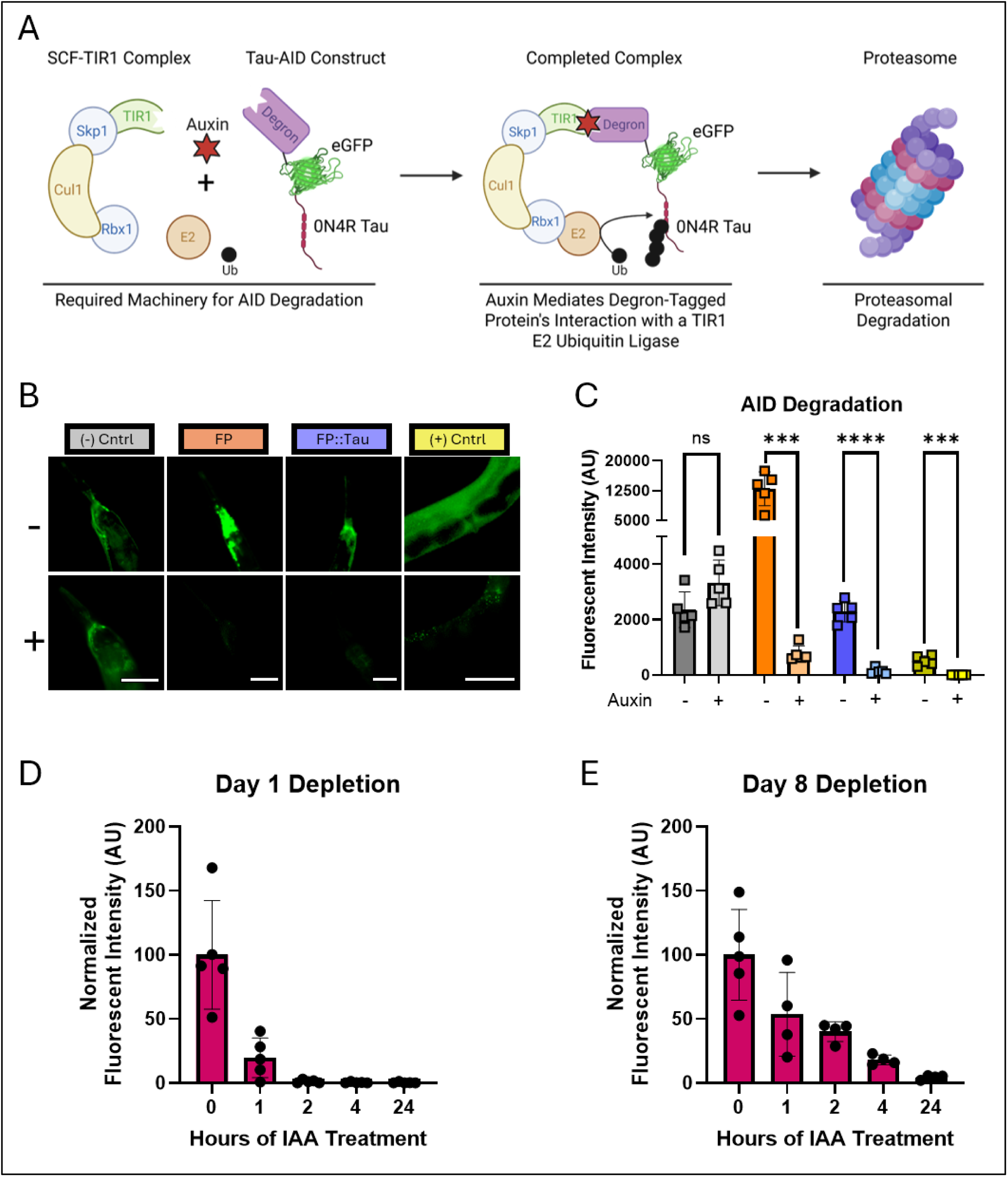
Auxin Treatment Rapidly Depletes Pan-Neuronal Tau (A) Schematic model illustrating the mechanism of auxin inducible depletion. Briefly, auxin mediates the interaction between the AID-tagged protein of interest and a TIR1-ubiquitin ligase complex, thereby targeting the tagged protein for proteasomal degradation. (B) Fluorescent images demonstrating FP::Tau levels without treatment (-) or with IAA auxin treatment for 24 hours (+). Scale bars are 50 µM. (C) Quantification of FP::Tau fluorescence with or without IAA treatment. Data shown is the backgrounded subtracted sum intensity of fluorescence in the head of each animal. Each bar represents the mean of each sample +/- SD with individual points representing individual animals. Individual t-test comparisons between untreated and IAA treated samples, p = 0.05, N = 5 animals. (D-E) Fluorescent quantification of auxin-mediated depletion of an integrated eGFP::T231E overexpression strain (KWN1025) at Days 1 (D) and 8 (E). Each point represents an individual animal, N = 4-5. For each time- course, fluorescent intensity was normalized to the starting intensity as a value of 100.

Because the single copy strains were quite dim (**Fig 1**), we utilized integrated overexpression strains to confirm rapid and complete depletion of the same AID-tagged tau transgene (**Fig 3B and 3C**). A pan- neuronal tau strain lacking the necessary cofactor TIR1 was used as a negative control, and a strain containing an AID tagged protein in smooth body wall muscle alongside pan-somatic TIR1 was used as a positive control (**Fig 3B**). To determine how quickly depletion occurred, we quantified fluorescent intensity over time following auxin exposure, in both young (Day 1) and aged (Day 8) animals. In Day 1 animals, approximately 99% of the fluorescently tagged tau had been degraded by 2 hours (**Fig 2D**).

While slower in Day 8 animals, 98% of the fluorescent protein was still degraded within 24 hours (**Fig 2E**). These results are consistent with previous reports using an AID system to degrade other targets and the known decrease in protein turnover seen in aged *C. elegans* ^36, 43^. The caveat here is that we are not able to ensure robust depletion in every neuron, particularly in the single-copy strains. However, this would only emerge as a concern if the depletion strategy failed to suppress the tau-induced phenotypic deficit.

Single-copy pan-neuronal T231E strains were crossed with the pan-neuronal TIR1 *otIs730* allele and assayed for touch responsiveness following auxin treatment. Assays were performed at adult Days 3, 5, and 8 to interrogate the role of age in the phenotypic deficit and its reversibility (**Fig 4A**). As expected, the touch deficit was exacerbated with age, but to our surprise, T231E depletion rescued the touch deficit completely at all ages tested, in a manner that was entirely dependent upon both TIR1 and IAA (**Fig 4B, 4C, and 4D**). This finding refutes our hypothesis that stress adaptation over time may result in irreversible abnormalities that preclude restoring normal neuronal function ^44^. We conclude that tau inhibition of mechanosensory neuronal function is a transient and completely reversible phenomenon, at least in our model, which bodes well for therapeutic clearance strategies’ effectiveness.

**Figure 4.**
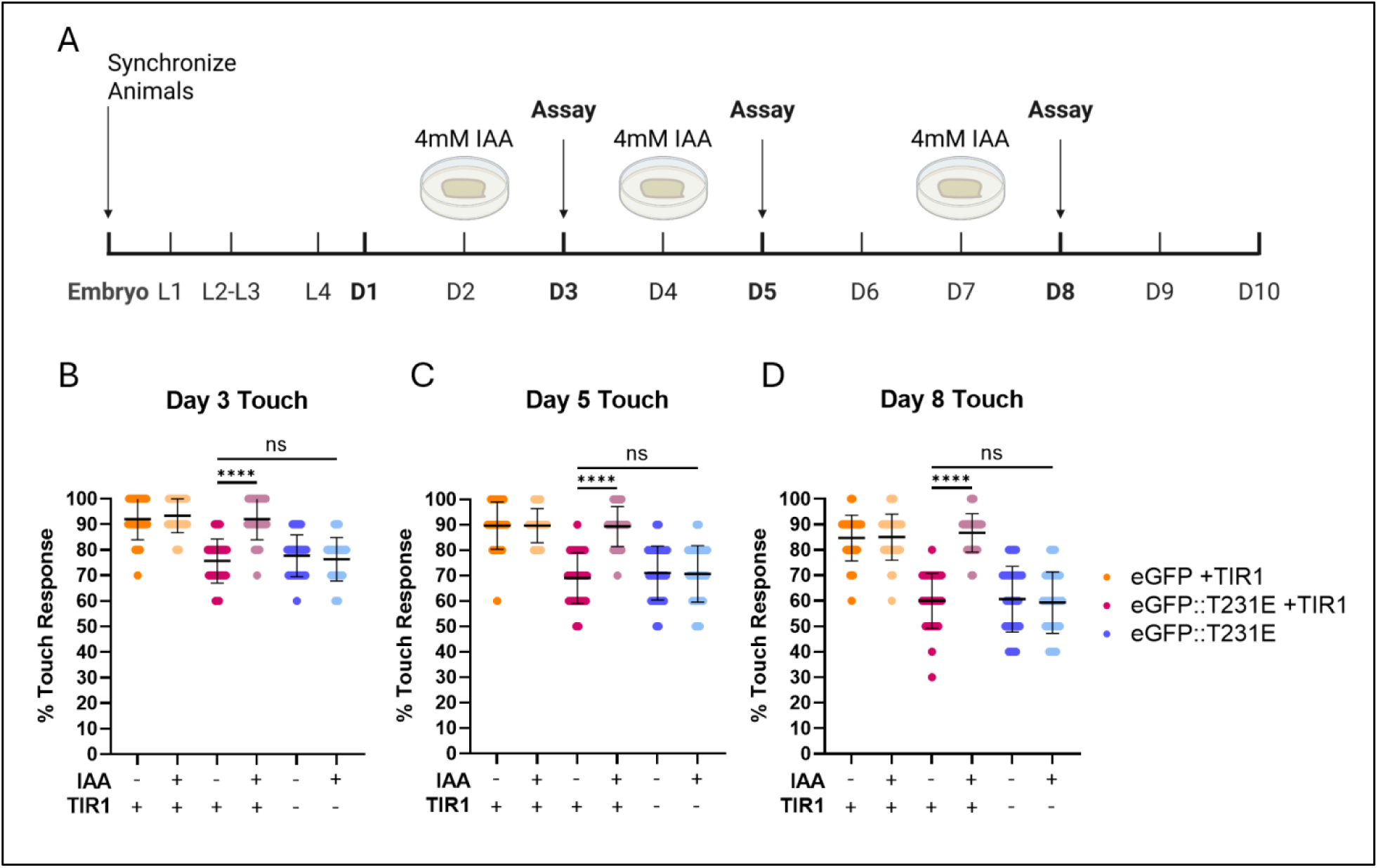
Tau Depletion via AID Rescues Touch Deficits at All Ages (A) A schematic demonstrating the experimental outline used to test how AID depletion of Tau affects eGFP::T231E-dependent touch deficits with age. Briefly, animals were synchronized and moved to control NGM plates or NGM plates containing 4mM IAA at adult days 2, 4, and 7, then assayed at days 3, 5, and 8, respectively. (B-D) Quantification of touch sensitivity at adult Days 3, 5, and 8. Pan-neuronal AID::eGFP strains containing pan-neuronal TIR1 were used as a negative control (red), and AID::eGFP::T231E strains lacking TIR1, and therefore incapable of tau depletion upon treatment with IAA, were used as a positive control (blue). Experimental strains containing both pan-neuronal AID::eGFP::T231E and TIR1 are denoted in yellow. Each dot represents an individual animal, N = 30 animals from three biological replicates for each condition. Statistical analysis was performed using a one-way ANOVA followed by Tukey-corrected multiple comparisons.

## Discussion

The AD field has largely acknowledged that tau’s primary toxicity likely occurs upstream of aggregation, as seminal studies have demonstrated that aggregates may be protective in disease contexts ^45, 46^. Many studies have demonstrated that specific tau phosphorylation events contribute to AD development ^47, 48^, yet the precise mechanisms through which these early-disease PTMS alter tau’s oligomerization and contribute to pathology is not clearly defined. A large contributor to this lack of information arises from the complexity and ambiguity inherent to disease models, particularly on those which rely on overexpression and report phenotypes which may not be relevant to AD pathology ^32, 33^. Modern approaches to AD diagnosis appreciate that molecular changes occur years before overt neurodegeneration and AD symptoms manifest ^49–51^. In this light, physiologic expression models which highlight subtle, yet reproducible phenotypes are of increasing importance, as they may help to illuminate the earliest events in tau pathogenesis.

Given the fact that the greatest risk factor for AD is aging, the field requires models which develop normally and possess normal lifespans, unlike overexpression models, which commonly have shortened lifespans and gross morphological differences ^32^. Our novel model expresses single copy levels of tau equally in all neurons using the UPN promoter (**Fig 1**) and possesses a mild, age-dependent touch deficit (**Fig 3**). The localization of both TauT4 and T231E tau to the neuronal processes suggests that the T231E mutation does not majorly impacting tau’s ability to bind microtubules (**Fig 1C**). The lifespan, stress resistance, and overall health of the animals seems unaffected by pan-neuronal T231E in any context tested (**Fig S1**). Furthermore, we are confident in the ability of the AID system to rapidly and robustly deplete our phosphomimetic tau at any age, given we observe nearly complete depletion of pan- neuronal AID-tagged proteins within 24 hours of 4mM IAA treatment at all ages (**Fig 2**).

Tau pathology in sporadic AD cases follows a defined spatial progression, indicating that certain regions of the brain are more susceptible to AD pathology ^52^. This phenomenon is broadly referred to as selective vulnerability, and understanding the mechanisms that confer susceptibility may provide insights into how early-disease tau modifications contribute to neuronal dysfunction. Selective vulnerability is not limited to brain regions, however, as specific neuronal subtypes in each region can alter susceptibility. For example, excitatory neurons in human AD cases are more prone to tau pathology than inhibitory neurons ^53^. Many studies have proposed mechanisms for this enhanced susceptibility, such as differential expression of tau isoforms ^54^ or differences in chaperone activity, such as observed with BAG3 ^53^.

As mentioned above, the mechanosensory neurons were the only neuronal subset that showed quantifiable deficits that could be attributed to cell autonomous action of tau, based on this work (**Fig 3A**) and previous publications ^15, 38^. Other commonly reported phenotypes observed in pan-neuronal tau models were not present, including thrashing deficits ^55^, overt locomotor deficits ^56^, and associative memory deficits ^41^. The specificity of the touch deficit in our model could indicate that the morphology of mechanosensory neurons plays a part in their vulnerability to tau pathology. Specifically, excitatory neurons typically possess much longer neuronal processes than inhibitory neurons ^53^. Similarly, the mechanosensory neurons in *C. elegans* possess the longest neuronal processes in the animal ^57^, indicating that axonal length or microtubule abundance may modulate AD susceptibility on a cellular scale. Single nucleus RNA sequencing within susceptible vs. resistant subpopulations of neurons within the entorhinal cortex Layer II (ECII) supports this finding by highlighting that vulnerable ECII neurons differentially express axon-localized proteins ^58^. Previously we demonstrated that axonal trafficking in mechanosensory neurons of T231E mutants is impaired and that mito-lysosomal trafficking along the mechanosensory neurite progressively declines with age ^38^. These observations highlight the intersection of tau, axonal maintenance, and mitochondrial trafficking as a critical component of incipient neuronal dysfunction in our AD model.

It is possible that simply clearing pathological proteins may not be sufficient for reversing disease progression, in part due to lasting consequences of the toxic moiety ^59^. The most obvious example of this would be if a toxic protein caused neuronal death, a phenomenon which protein clearance could not reverse in a population of post-mitotic neurons. Permanent changes could also occur in more subtle ways, such as through maladaptive stress responses. For example, we hypothesized that tau may cause irreversible metabolic changes in neurons by triggering chronic mitochondrial stress and eliciting the mitochondrial unfolded protein response. Several studies, including our own ^15, 38, 60, 61^, have demonstrated that tau containing disease-associated PTMs can affect mitochondrial function and mitochondrial quality control (MQC), a set of dedicated molecular pathways designed to ensure healthy mitochondrial networks ^60–62^. Chronic mitochondrial unfolded protein response signaling can inhibit mitophagy, a form of selective autophagy specific to mitochondria and a key part of MQC ^63^. Long-term disruption of MQC skews mitochondrial homeostasis through the accumulation of damaged mitochondria and abnormal mitochondrial DNA ^63, 64^. Maladaptive changes such as these present a potential confound for studies and therapies based on the protective effects of tau reduction and clearance ^65–67^, which may only be effective if administered before adaptive responses become chronic. However, despite our hypothesis that T231E would cause irreversible changes as animals aged, we saw a complete rescue of the touch deficit regardless of the age at which tau was depleted (Figure 4).

While the touch assay measures the direct function of mechanosensory neurons, we also observed that the pan-neuronal T231E animals have discernable alterations in their curvature as they move on solid media (**Fig 3B and 3E**). Coordinated movement in *C. elegans* relies on a complex network of excitatory and inhibitory neurons ^68^, and T231E’s impact on curvature suggests that perhaps synaptic signaling throughout the neuronal motor circuit is impaired. Further studies in our laboratory aim to elucidate if specific neuronal subsets within this circuit are selectively impacted, or whether synaptic transmission is less efficient within this model. Additionally, the subtle differences in T231E tau animal’s curvature are likely the earliest manifestation of overt locomotor deficits observed in other tau overexpression models ^32^. This further highlights the importance of carefully accounting for tau expression levels in animal models to separate physiologically relevant disease findings from artifacts of protein overexpression. In sum, our results suggest that the pathologic effect of T231E is completely reversible, but they also highlight that early-disease phosphorylation events such as occur at T231 might independently lead to subtle, pre-clinical neuronal dysfunction.

Most clinical trials for tau clearance are currently in the early stages. Many of these are based on immunotherapy and rely on antibodies to target tau, with degradation occurring through the host immune system ^67^. Our results suggest that therapies which degrade tau species associated with early-disease progression can prevent subtle neuronal dysfunction and may protect neurons from stress that could culminate into more severe neuronal damage with age. Unlike our AID approach, however, many Alzheimer’s clearance strategies target extracellular tau, which have limited intracellular access and thus may slow tau’s transfer between neurons but ultimately fail to prevent disease ^69^. In this light, the most effective therapies will likely be those which target a gamut of early-disease tau epitopes and degrade them before toxic tau incites intracellular stress and before more overt symptoms manifest ^70^.

## Conclusion

Our study uses a single-copy approach in *C. elegans* to model the functional impact of pan-neuronal tau expression, focused on achieving very low tau expression levels and equivalent expression in each neuron. Additionally, we tested whether phenotypic defects were reversible via tau depletion using an AID approach. This is particularly relevant to AD in older worms where irreversible changes or maladaptation may have occurred through life-long exposure to toxic tau. We demonstrated surprising selectivity in the behavior(s) impacted by an AD-relevant phosphomimetic tau, which is one of the earliest biomarkers of disease. Finally, we clearly showed that neuronal dysfunction caused by phosphomimetic human tau is entirely reversible, regardless of age, supporting the idea that tau clearance may prove an effective therapy for slowing AD progression. Further study is needed to determine the molecular mechanisms behind the selective and transient mechanosensory dysfunction observed in our model, and connecting single-copy to overexpression phenotypes will benefit from models that express defined amounts of tau from safe-harbor insertions such as described here that bridge a range of expression levels from low to high.

## Acknowledgements

The authors would like to thank the lab of Oliver Hobert for providing plasmids containing the UPN promoter. We would also like to acknowledge BioRender for aide in figure preparation. Additionally, we would like to thank Dylan Pfendler for helpful discussions during this work.

## Author Contributions

Carroll, T. - Conceptualization, Data Curation, Writing – Original Draft, Writing – Review and Editing, Visualization. Nehrke, K. and Johnson, G.V.W. – Writing – Review and Editing, Supervision.

## Ethical Considerations

Not applicable. **Consent to Participate** Not applicable.

## Consent for Publication

Not applicable.

## Data Availability Statement

The data supporting the findings of this study are available within the article and/or its supplemental material.

---

**Supplemental Figure S1.**
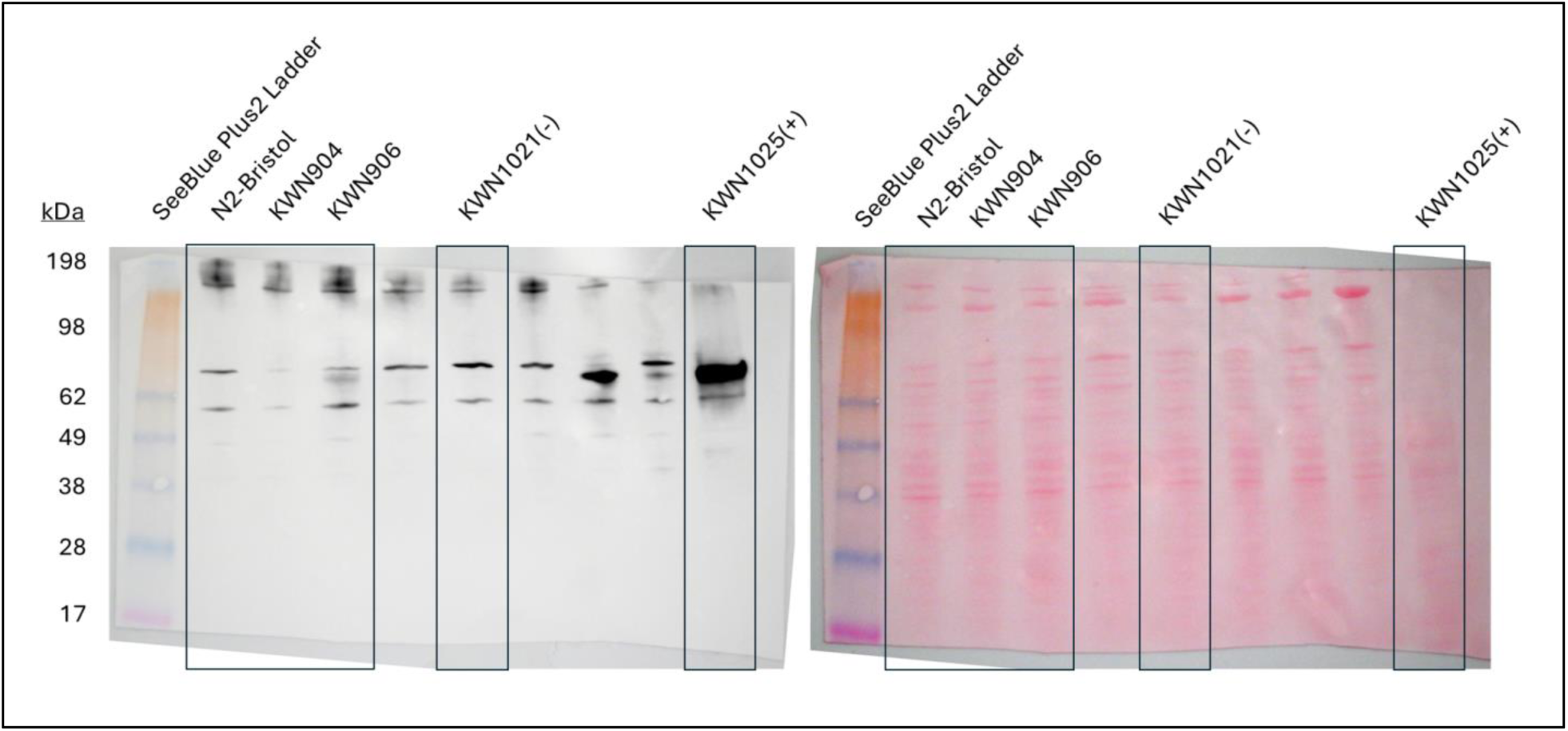
Full Lane Overview of Single-Copy Tau Western Blot Full-scale images of the Western Blot from Fig. 1A. Lanes shown in Fig. 1A are denoted by black boxes. Lysates from N2, single-copy eGFP (KWN904), single-copy T231E (KWN906), a strain overexpressing eGFP (KWN1021) as a negative control, and a strain overexpressing T231E (KWN1025) as a positive control were separated on a 4-15% gradient polyacrylamide gel and transferred to a nitrocellulose membrane. All samples were run on the same gel, transferred to the same membrane, and imaged with the same exposure time. Ponceau staining (right) was used to qualitatively visualize protein content in each lane. The membrane was probed using a rabbit, anti-tau antibody (1° - A0024, Dako, 1:150,000) overnight, washed three times using TBS-T, and then probed using a horse radish peroxidase (HRP) conjugated anti-rabbit antibody (2° - R1006, Kindle Biosciences, 1:5000) for 1 hour.

**Supplemental Figure S2.**
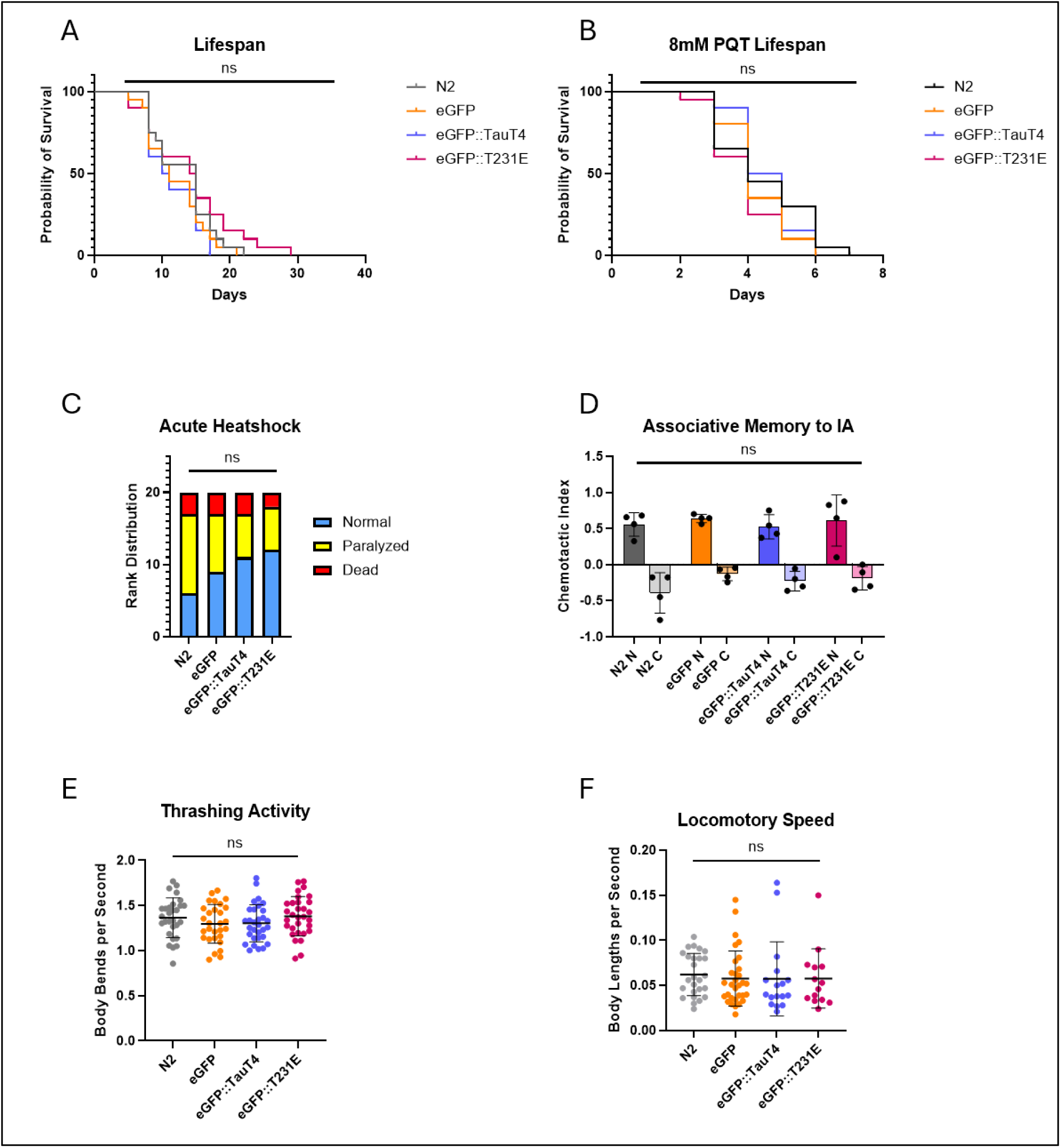
Phenotypes Not Impacted by Tau (A) Survival curve demonstrating baseline lifespan of the single-copy Tau mutants versus controls. N = 1 replicate of 20 individual animals. (B) Survival curve during chronic exposure to 8mM PQT. N = 1 replicate of 20 individual animals. (C) Score distribution of Tau expressing Day 1 animals after 1.25 hours of heat shock at 37°C. Animals were categorized as dead if no movement was observed after gentle prodding with a platinum wire, and they were considered paralyzed if they could move their head, but not the rest of their body. N = 1 replicate of 20 individual animals. (D) Chemotactic indexes to isoamyl alcohol (IA), normally an attractant for *C. elegans*, reflecting the associative memory when animals are exposed to IA in the absence of food. “N” denotes the non-conditioned starvation control for each strain, whereas “C” denotes the starvation + IA conditioned sample. For each strain, animals were initially attracted to IA but learned to avoid it when starved in its presence. Each point represents a population of 50-100 animals, N=4. (E) Quantification of thrashing rate at Day 5 in M9 liquid after 1 minute of acclimatization. Each point represents an individual animal. F) Normalized quantification for the average speed of Day 5 animals in body lengths per second. Each point represents an individual animal.

